# KCC2 is required for the survival of mature neurons but not for their development

**DOI:** 10.1101/2020.10.29.360875

**Authors:** Georgina Kontou, Josephine Ng, Ross Andrew Cardarelli, Jack Howden, Catherine Choi, Qiu Ren, Josef Kittler, Nicholas Brandon, Stephen J. Moss, Joshua L. Smalley

**Affiliations:** AstraZeneca-Tufts Laboratory of Basic and Translational Neuroscience, Tufts University School of Medicine, 136 Harrison Ave, Boston, MA 02111; Department of Neuroscience, Tufts University School of Medicine, 136 Harrison Ave, Boston, MA 02111; Department of Neuroscience, Physiology, and Pharmacology, University College London, WC1E 6BT, UK; Neuroscience, IMED Biotech Unit, AstraZeneca, Boston, MA 02451

**Keywords:** apoptosis, extrinsic pathway, cell death, seizures

## Abstract

The K+/Cl– co-transporter KCC2 (SLC12A5) allows mature neurons in the CNS to maintain low intracellular Cl^−^ levels that are critical in mediating fast hyperpolarizing synaptic inhibition via type A γ-aminobutyric acid receptors GABA_A_Rs. In accordance with this, compromised KCC2 activity results in seizures but whether such deficits directly contribute to the subsequent changes in neuronal viability that lead to epileptogenesis, remains to be assessed. Canonical hyperpolarizing GABA_A_R currents develop postnatally which reflect a progressive increase in KCC2 expression levels and activity. To investigate the role that KCC2 plays in regulating neuronal viability and architecture we have conditionally ablated KCC2 expression in developing and mature neurons. Decreasing KCC2 expression resulted in the rapid activation of the extrinsic apoptotic pathway, in mature hippocampal neurons. Intriguingly, direct pharmacological inhibition of KCC2 in mature neurons resulted in the rapid activation of the extrinsic apoptotic pathway. In contrast, ablating KCC2 expression in immature neurons had no discernable effects on their subsequent development, arborization or dendritic structure. However ablating KCC2 expression in immature neurons was sufficient to prevent the subsequent postnatal development of hyperpolarizing GABA_A_R currents. Collectively, our results demonstrate that KCC2 plays a critical role in neuronal survival by limiting apoptosis, and mature neurons are highly sensitive to loss of KCC2 function. In contrast KCC2 appears to play a minimal role in mediating neuronal development or architecture.

## INTRODUCTION

The K^+^/Cl^−^ co-transporter KCC2 (encoded by the gene *SLC12A5*) is the principal Cl^−^-extrusion mechanism employed by mature neurons in the CNS (1). Accordingly, its activity is an essential determinant for the efficacy of fast synaptic inhibition mediated by type A γ-aminobutyric acid receptors (GABA_A_R), which are Cl^−^ permeable ligand-gated ion channels. At prenatal and early postnatal stages in rodents, neurons have elevated intracellular Cl^−^ levels resulting in depolarizing GABAAR-mediated currents (2). The postnatal development of canonical hyperpolarizing GABA_A_R currents is a reflection of the progressive decrease of intraneuronal Cl^−^ that is caused by the upregulation of KCC2 expression and activity, which do not reach their maximal levels in humans until 20-25 years of age (3). Consistent with this critical role in regulating inhibitory neurotransmission, patients with mutations in *SLC12A5* exhibit severe epilepsy that develops after birth, together with severe developmental delay (4–6). The essential role KCC2 plays in the brain is further exemplified by gene knock-out studies in mice where homozygotes die shortly after birth (7, 8). In addition to neuronal Cl^−^ homeostasis, KCC2 has been proposed to play a role in determining the morphology of dendritic spines that is independent of its Cl^−^ transport function (9–11).

Alterations in KCC2 function have been reported in patients with temporal lobe epilepsy (TLE) (12–14). In addition to spontaneous seizures, patients with TLE have profound changes in the structure of the hippocampus which includes extensive neuronal death, modified arborization of surviving neurons, and reactive gliosis. Significantly, we have recently shown that local inactivation of KCC2 in the hippocampus is sufficient to induce the development of spontaneous seizures resembling those seen in TLE (15). However, the direct contribution of KCC2 expression levels and/or activity to the neuronal loss and anatomical changes seen in TLE are unknown.

To investigate the role that KCC2 plays in determining neuronal viability, we have examined the consequences of ablating its expression prior to and following neuronal development. In mature cultured neurons and the adult hippocampus, decreasing KCC2 expression activated the extrinsic pathway of apoptosis resulting in rapid neuronal death. Acute pharmacological blockade of KCC2 also induced apoptosis by a similar mechanism. Ablating KCC2 expression levels in immature neurons compromised the development of hyperpolarzing GABA_A_R currents but did not impact on neuronal viability, proximal dendritic arborization or spine formation. Collectively our results suggest that KCC2 is required for survival of mature neurons and hyperpolarizing GABA_A_R currents but is dispensable for their development and morphology.

## RESULTS

### Cre-dependent ablation of KCC2 expression in mature and immature neurons

In order to investigate the significance of KCC2 for neuronal development, maintenance, and viability, we generated hippocampal cultures from P0 mice in which *loxP* sites have been introduced into the *SLC12A5* gene (KCC2^fl^). These cultures were then infected with adeno-associated viruses; *AAV9-CaMKII-eGFP* (AAV-GFP), or *AAV9-CaMKII-eGFP-Cre* (AAV-Cre). Using the CaMKII promoter, we were able to target Cre expression to principal neurons after 3 (immature) or 18 (mature) days *in vitro* (DIV) (**Figure 1a-c**). The expression of Cre recombinase leads to the excision of exons 22-25 between the two *loxP* sites inserted into the *SLC12A5* gene (**Figure 1a**) resulting in a translated protein that lacks amino acids Q911-Q1096 of KCC2 that are essential for transporter activity (15, 16) (**Figure 1b**).

**Figure 1.**
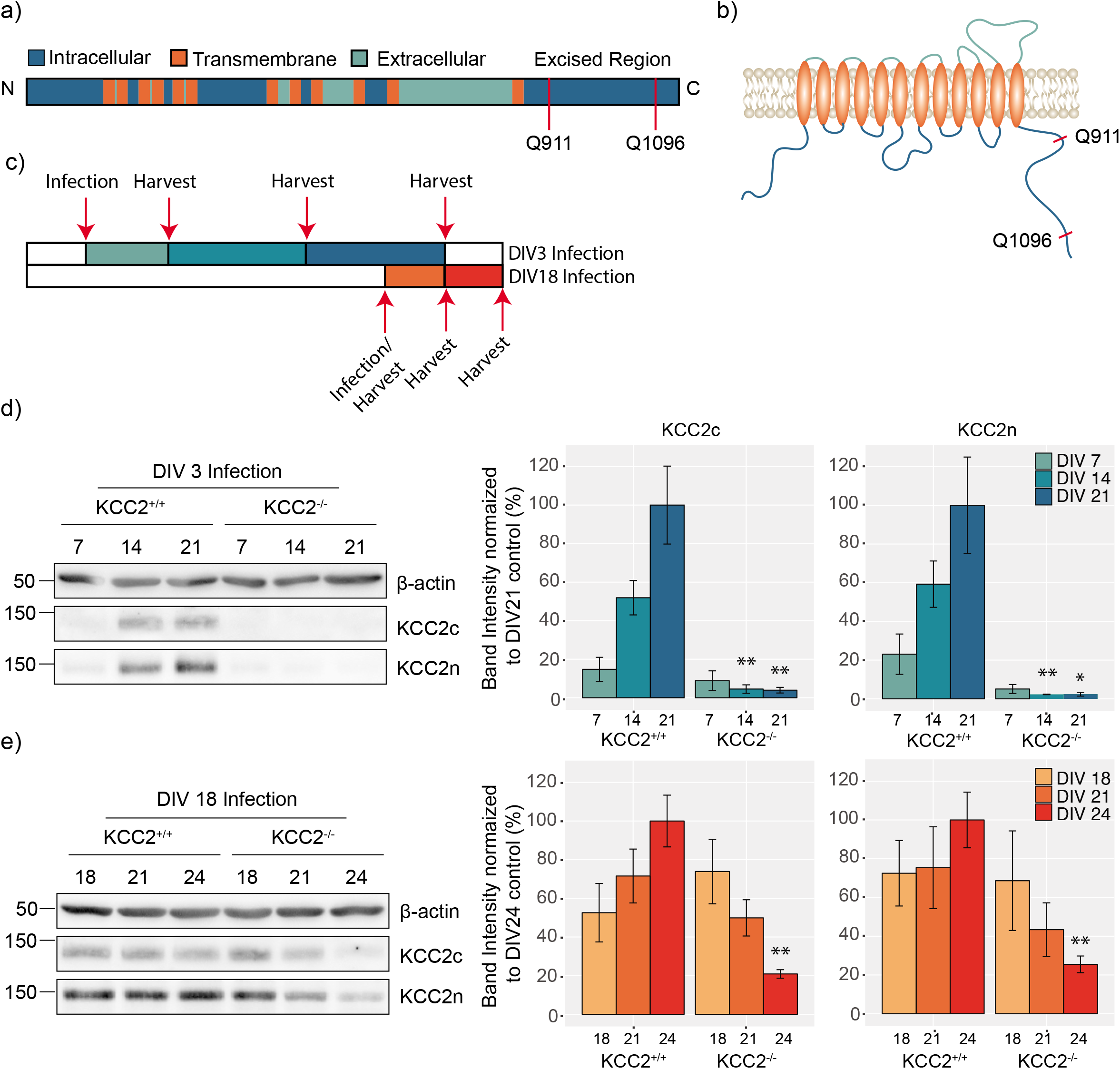
Using a viral approach to knock-out KCC2 from mature and developing dissociated hippocampal neurons. **A.** Schematic diagram of the KCC2 gene, denoting the region excised by the Cre-recombinase. **B.** Schematic diagram of KCC2 protein topology, denoting the excised region on the intracellular C-terminus. **C.** Schematic diagram of viral treatment paradigm to knock-out KCC2 in developing (DIV3) and mature (DIV18) neuronal networks. **D.** Immunoblots and corresponding bar chart quantification confirming the reduction of KCC2 expression after AAV-Cre viral infection of KCC2^fl^ hippocampal dissociated neurons at DIV3. **E.** Immunoblots and corresponding bar chart quantification confirming the reduction of KCC2 expression after AAV-Cre viral infection of KCC2^fl^ hippocampal dissociated neurons at DIV18.

To determine the efficiency of recombination, immature neurons infected at DIV3 were harvested at DIV7, DIV14, and DIV21, lysed and immunoblotted with KCC2 antibodies (**Figure 1c**). Using a C-terminal antibody, KCC2 expression levels increased over this time course in neurons infected with AAV-GFP. In neurons infected with AAV-Cre, KCC2 levels were reduced to 4.5 ± 2.18% and 3.9 ± 1.46% at DIV14 and 21 respectively compared to control (p_DIVI4_ = 0.007, p_DIV21_ = 0.009) (**Figure 1d**). Using an N-terminal antibody, KCC2 expression levels were also reduced significantly in cultures infected with AAV-Cre (DIV14: 2.3 ± 0.14%, *p* = 0.009, DIV21: 2.4 ± 0.99%, *p* = 0.049). To assess the efficiency of KCC2 removal in mature networks, we harvested neurons for western blot analysis on the day of (DIV18), 3 days (DIV21) and 6 days (DIV24) after viral infection (**Figure 1c**). The total protein levels of KCC2 were gradually reduced 3 and 6 days after infection (**Figure 1e**). By DIV24, KCC2 levels were reduced to 20.9 ± 2.21% (*p* = 0.004) and 25.5 ± 4.30% (*p* = 0.008) in the AAV-Cre compared to the control, as detected by the C- and N-targeting antibodies respectively. Collectively these results demonstrate that viral infection using AAV-Cre is an efficient strategy to reduce KCC2 expression levels in developing and mature neuronal networks.

### Reducing KCC2 expression does not modify neuronal arborization proximal to the soma

To assess the effects of reduced KCC2 expression on neuronal morphology we employed immunofluorescence (**Figure 2a**). Firstly, we measured KCC2 expression levels. KCC2 expression was significantly decreased in neurons infected with AAV-Cre, which were identified by the presence of GFP. We observed a 63.4 ± 2.27% and a 54.6 ± 2.50% decrease in the KCC2 fluorescence levels in GFP-positive regions in developing and mature neuronal networks respectively in the AAV-Cre compared to the AAV-GFP control (**Figure 2b,** *P*_DIV3_ < 0.001, *P*_DIV18_ < 0.001).

**Figure 2.**
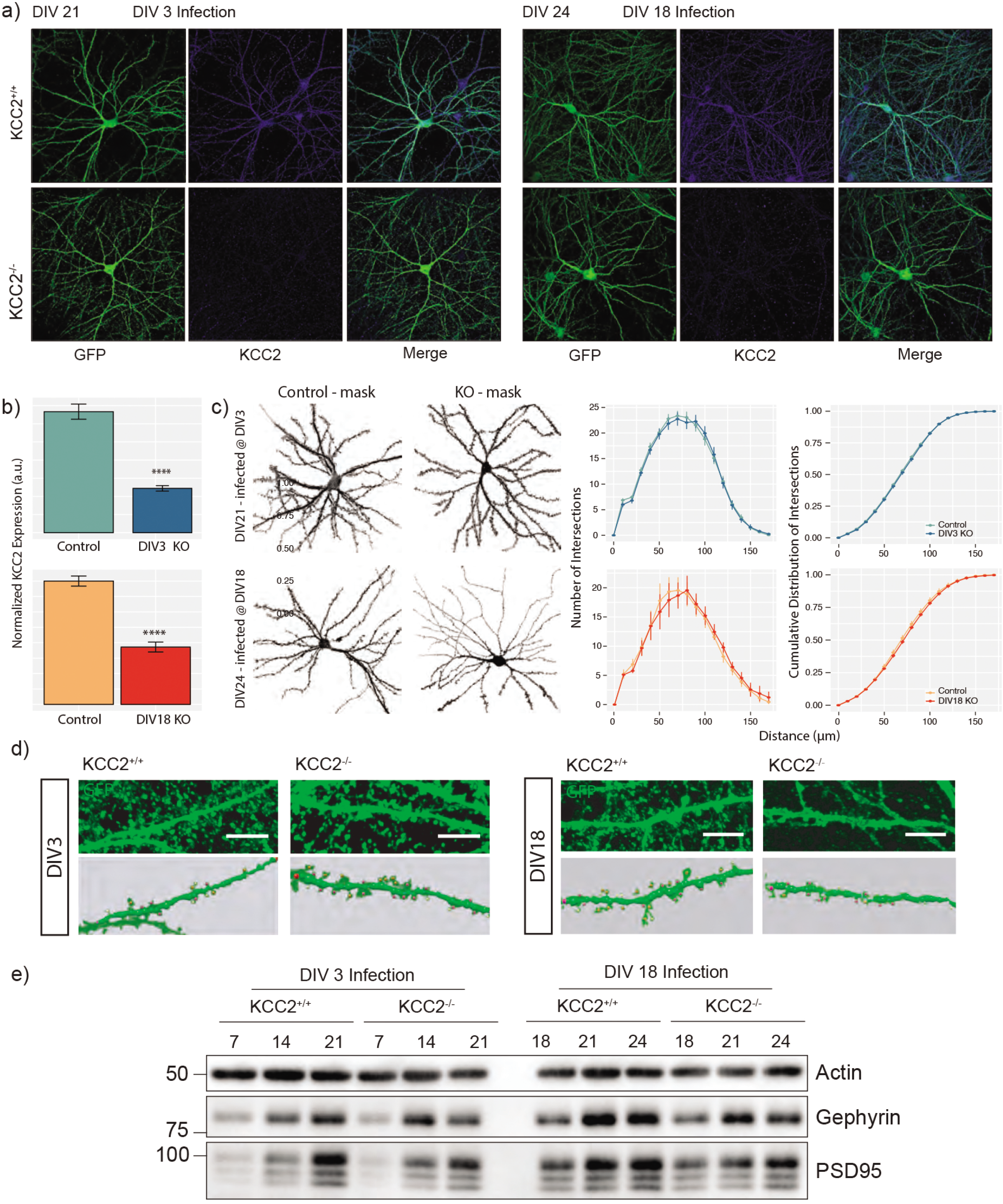
The effect of removing KCC2 from developing and mature networks on maintenance of dendrites, spines and synapses. **A.** Primary cultured neurons from P0 pups were infected with AAV-GFP or AAV-Cre at DIV3 and DIV18 and fixed at DIV21 and DIV24 respectively. The cells were immunostained for KCC2 and GFP to amplify the signal, and confocal images were acquired using a 60X objective (Scale bar = 10 μm). **B**. Quantification of the normalized mean fluorescent KCC2 signal within GFP positive neurons for DIV3-infected and DIV18-infected cells. **C.** Neuronal Reconstruction of control and knock-out neurons infected with the AAV-GFP or AAV-Cre for Sholl Analysis using Simple Neurite Tracer Fiji plugin. The number of intersections (y-axis) are plotted against distance from the soma (x-axis) and as a cumulative distribution. **D.** Cropped confocal images of GFP-expressing neuronal processes with dendritic spines (Scale Bar = 10 μm) with spine reconstructions below, made using Neuron Studio (40). **E.** Immunoblots showing the levels of the excitatory postsynaptic marker PSD95 and the inhibitory postsynaptic marker gephyrin in AAV-Control and AAV-Cre infected cells at DIV3 and DIV18.

The expression of the GFP fluorophore allowed us to reconstruct the neuronal morphology of CaMKII-positive pyramidal cells in the presence and absence of KCC2 (**Figure 2c**). Using Sholl analysis, it was apparent that reducing KCC2 expression did not modify the complexity of proximal dendrites close to the cell body, as there was no morphological difference from the control in either experimental groups (**Figure 2c**). Likewise, the number of intersections between the dendritic segments were also unaffected at either time point.

We also tested if reduced KCC2 expression impacted on the formation of dendritic spines. The loss of KCC2 had no robust effect on dendritic spines, as spines were still present in the absence of KCC2 from both developing and mature networks (**Figure 2d**). Interestingly, an increase in stubby structures was observed when KCC2 was removed from mature networks which could reflect an increase in immature spines. Consistent with this, a modest reduction in both gephyrin and PSD95 (inhibitory and excitatory post-synaptic markers respectively) was evident in mature neurons deficient in KCC2 (**Figure 2e**). Collectively these results suggest that KCC2 does not play a major role in regulating neuronal arborization or dendritic structure.

### Ablating KCC2 expression prevents the development of hyperpolarizing GABA_A_R currents

We performed gramicidin perforated-patch recordings on AAV-GFP or AAV-Cre infected neurons to examine the effects of reducing KCC2 expression levels on the reversal potential of GABA-induced currents (E_GABA_), which in mature neurons is predominantly set by KCC2 activity. E_GABA_ was measured by calculating the reversal potential of leak-subtracted muscimol currents (1 μM) during positive-directed voltage ramps. To isolate the contribution of KCC2 to E_GABA_, the NKCC1 inhibitor bumetanide (10 μM) was applied and neuronal activity was blocked using tetrodotoxin (TTX; 500 nM) to prevent activity-dependent Cl^−^ accumulation. Removing KCC2 from developing networks resulted in more depolarized E_GABA_ values in the AAV-Cre infected cells compared to AAV-GFP infected controls, demonstrating the critical role of the KCC2 transporter (**Figure 3c,** Control_DIV3_: −106.7 ± 2.42 mV, Cre_DIV3_: −64.2 ± 2.52 mV, *p*< 0.001). This was also reflected as an increase in the intracellular concentrations of Cl^−^ when KCC2 was removed (**Figure 3d,** Control_DIV3_: 2.4 ± 0.22 mM, Cre_DIV3_: 12.7 ± 1.21 mM, *p*< 0.001). Furthermore, treatment with the selective KCC2 inhibitor, VU0463271 (11K; 1 μM), led to minimal shifts in E_GABA_ values in AAV-Cre infected cells, supporting the absence of KCC2 (**Figure 3e,** Control_DIV3_: 44.7 ± 3.25 mV, Cre_DIV3_: −3.2 ± 2.69 mV, *p* < 0.001). In summary these results are consistent with the well supported hypothesis that KCC2 expression is critical for the postnatal development of hyperpolarizing GABA_A_R currents.

**Figure 3.**
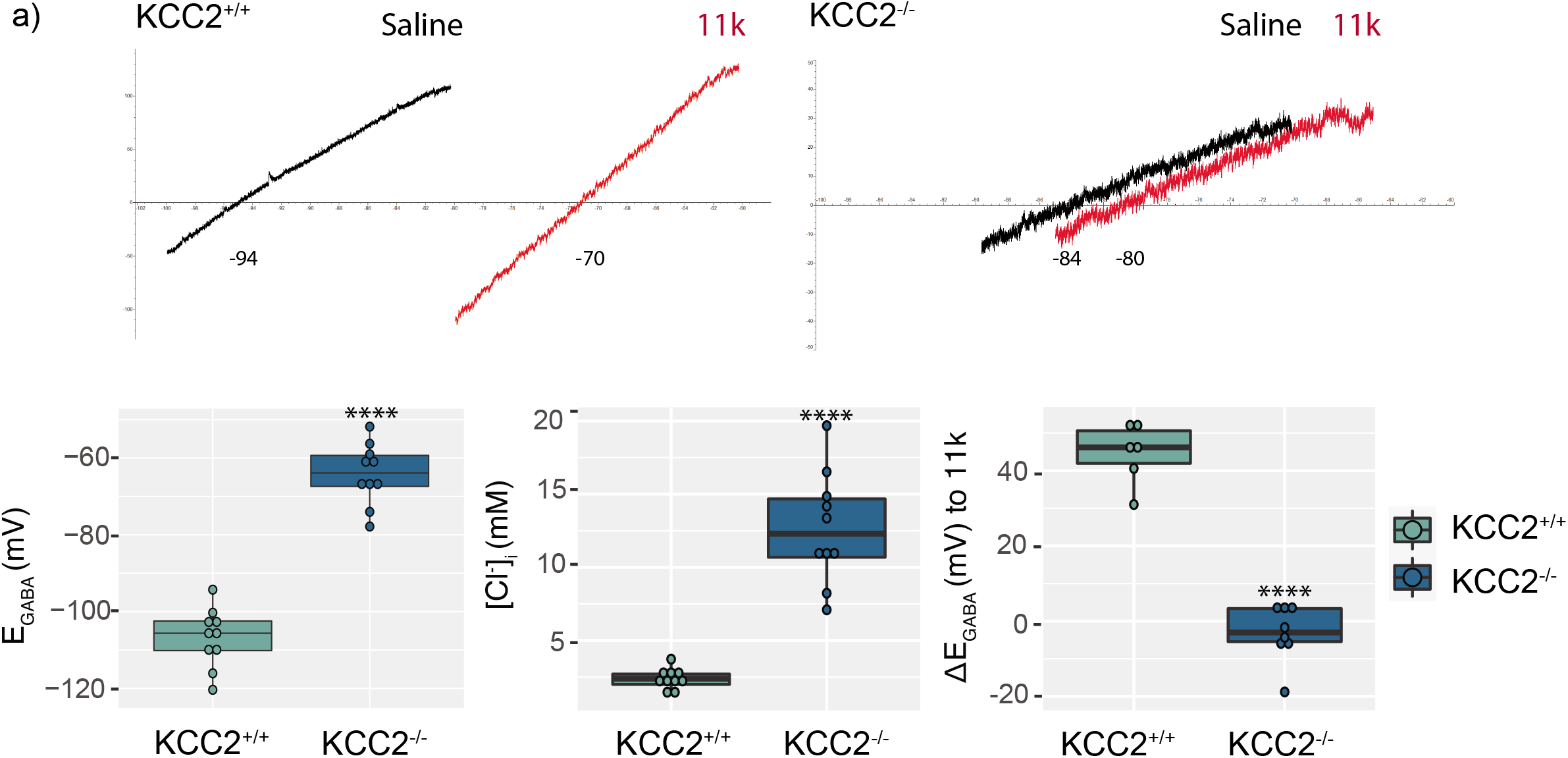
The effect of KCC2 knock-out on E_GABA_ and intracellular Cl^−^levels. **A.** Representative electrophysiological traces of DIV21 neurons infected with AAV-GFP or AAV-Cre at DIV3 showing the leak-subtracted muscimol-activated currents and the shift after 11K application. The box plots show the quantification for the mean E_GABA_, calculated Cl^−^ concentration and 11K-induced shift.

### Inactivation of KCC2 in mature neurons induces apoptosis

0Previous work from our lab has shown that a reduction of KCC2 levels decreases the viability of neuronal cultures (15). In order to understand the role of KCC2 and deregulated Cl^−^ homeostasis in the modulation of neuronal viability, we performed western blot analysis and probed neuronal lysates with known apoptotic markers, after viral infection with AAV-GFP or AAV-Cre (**Figure 4**). Under control conditions, the levels of cleaved caspases were low when cells were treated with AAV-GFP at both infection time points (DIV3 and DIV18). In contrast, a 90.1 ± 31.78% increase in the signal of the early proapoptotic marker; cleaved caspase 8, was evident at DIV21, when KCC2 was knocked-out at DIV18 (**Figure 4,** *p* = 0.047). The increase in cleaved caspase 8 coincides with a loss of KCC2 signal in mature neurons infected with the AAV-Cre, compared to the AAV-GFP infected control (**Figure 1, 4).** Interestingly, there were no changes in the cleavage of caspases associated with intrinsic pathway activation (caspase 9) when KCC2 was knocked-out from either immature or mature neurons. In parallel, a 58.4 ± 18.14% and a 37.3 ± 5.70% increase in the levels of the late proapoptotic markers; cleaved caspase 3, and cleaved poly (ADP-ribose) polymerase (PARP) respectively, were seen by DIV24 (**Figure 4,** *p*cl.caspase3 0.032, *p*cl.PARP 0.023). Thus, the sustained absence of KCC2 by DIV24 resulted in increased levels of active apoptotic executioner caspases, that lead to cell death via the activation of the extrinsic apoptotic pathway.

**Figure 4.**
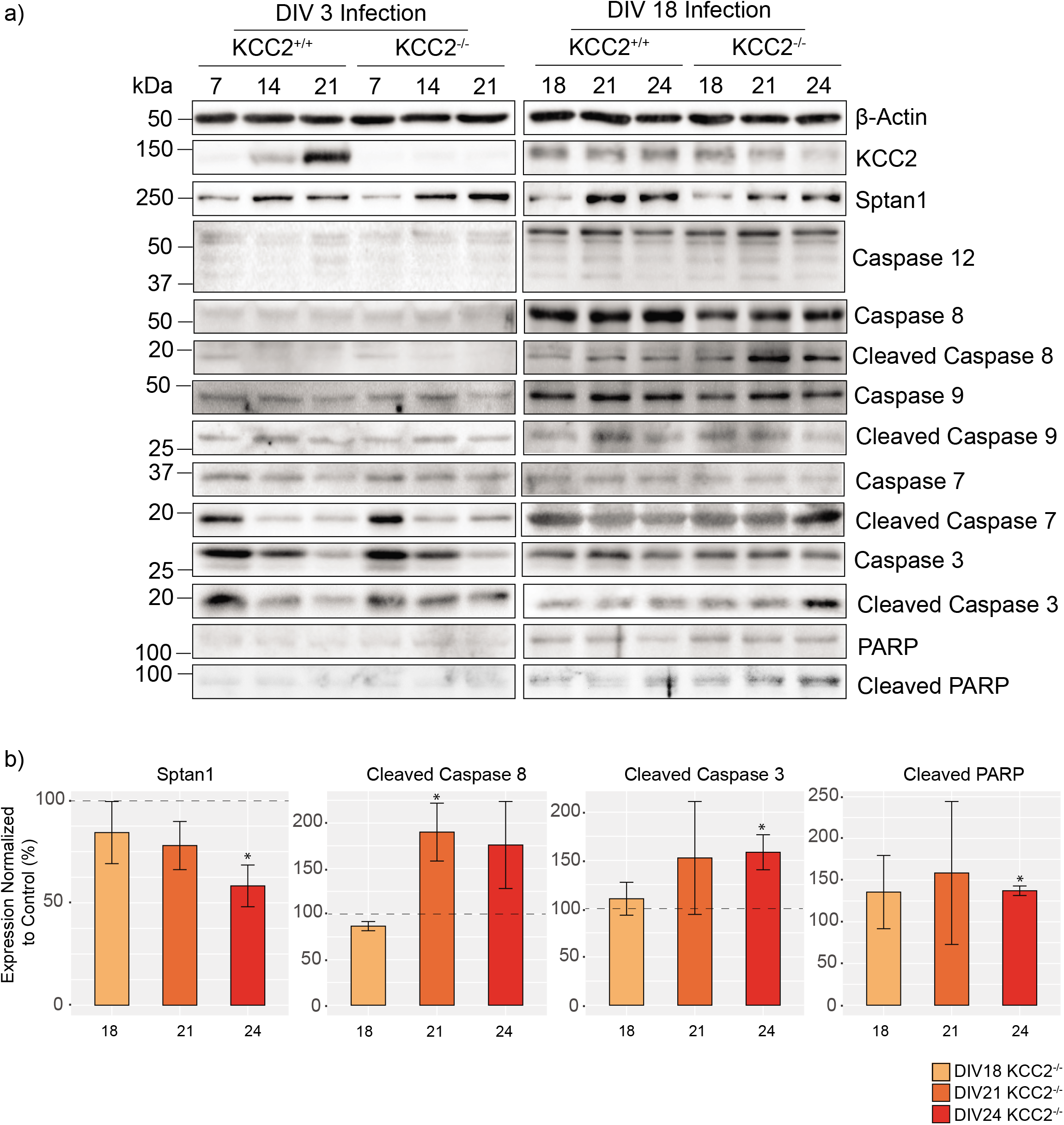
Reduced KCC2 expression in mature, but not developing neuronal networks increases the levels of proapoptotic cleaved caspases. **A.** Immunoblots on lysates of hippocampal cultures infected with AAV-GFP or AAV-Cre at DIV3 or DIV18. Lysates were probed for the total and cleaved levels of the extrinsic proapoptotic marker; caspase 8, the intrinsic proapoptotic markers; caspase 3 and caspase 7, the inflammation proapoptotic caspase 12, PARP, KCC2, the KCC2-associated structural protein Sptan1 and β-actin. **B.** Removal of KCC2 from mature networks resulted in an increase in the expression of several apoptotic markers. The bar charts show the quantification of the levels of cleaved caspase 8, cleaved caspase 3, cleaved PARP and Sptan1, with significantly changed expression marked (\**p* < 0.05).

In the brain, KCC2 is associated with Sptan1 in large macromolecular complexes (17). The levels of Sptan1, a previously characterized calpain and caspase 3 substrate and marker of neuronal damage (18, 19), were decreased to 58.3 ± 10.20% by DIV24 in the absence of KCC2 (**Figure 4,** *p* = 0.015). In contrast, there were no changes in the levels of apoptotic markers in cells infected with AAV-Cre at DIV3, suggesting that KCC2 removal from developing networks does not impact on neuronal viability. Thus, our results suggest that in mature but not developing neuronal networks, loss of KCC2 induces apoptotic cell death.

### Acute pharmacological inhibition of KCC2 induces apoptosis

To investigate the role that KCC2 activity plays as a determinant of neuronal viability further, we examined the effects of its acute inhibition on apoptotic markers. To do so, DIV18-21 cultured neurons were exposed to the selective KCC2 inhibitor 11K or DMSO (control) for 10, 30 and 60 min, and subsequently lysed and subjected to immunoblotting (**Figure 5**). Exposure to 11K increased the level of the early apoptotic marker; cleaved caspase 8 by 89.4 ± 5.98% compared to the control, within 10 min (*p* = 0.004). This was followed at 30 min by a 72.5 ± 23.78% increase in the late apoptotic marker; cleaved caspase 3 (*p* = 0.038). Furthermore, we also observed a striking 4-fold increase in the levels of cleaved PARP (*p* = 0.006). Thus, inhibition of KCC2 is sufficient to activate the apoptotic cascade.

**Figure 5.**
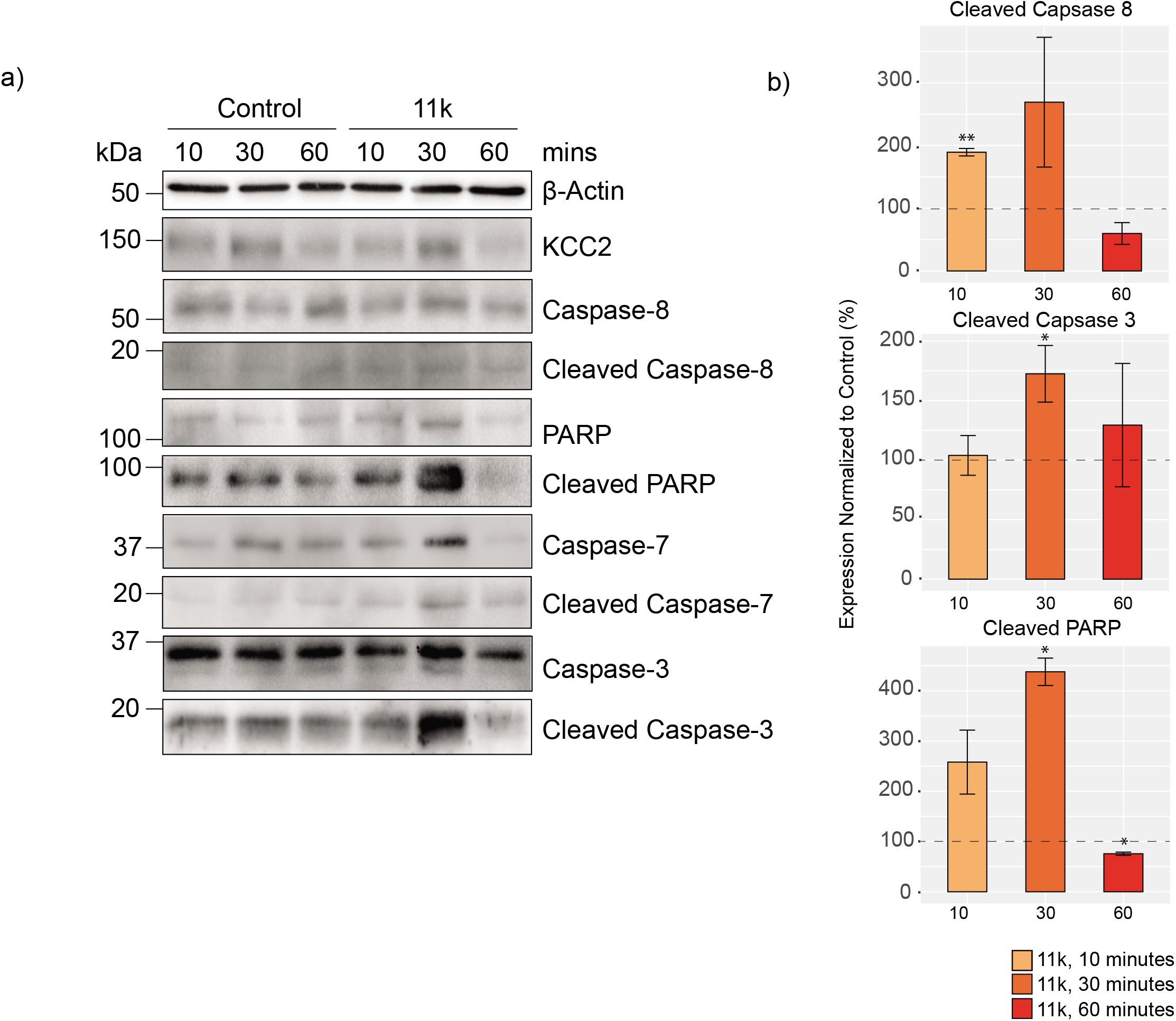
Pharmacological inhibition of KCC2 with 11K stimulates apoptosis in neuronal cultures. **A.** Immunoblots of DIV18-DIV21 hippocampal neuronal lysates treated with DMSO or 11K for 10, 30 or 60 minutes were probed for the total and cleaved levels of the extrinsic proapoptotic marker; caspase 8, the intrinsic proapoptotic markers; caspase 3 and caspase 7, PARP, KCC2, and β-actin. **B.** The bar charts show the quantification of the levels of proapoptotic markers cleaved caspase 8, cleaved caspase 3 and cleaved PARP, with significantly changed expression marked (\**p* < 0.05, \*\**p* < 0.01).

### Local inactivation of KCC2 in the hippocampus induces apoptosis

To assess the significance of our results in cultured neurons, we looked at the effects of reducing KCC2 expression on apoptotic markers in the brain. To do so, stereotaxic injections were used to introduce AAV-GFP or AAV-Cre, into the hippocampus of KCC2^fl^ mice. 7, 14 and 21 days following injection, immunoblotting was used to measure the levels of apoptotic markers between treatment groups from microdissected tissue identified by GFP expression (**Figure 6**). Consistent with previous data, infection with the AAV-Cre was sufficient to reduce the expression of KCC2 in mouse hippocampal neurons. The total KCC2 levels dropped to 59 ± 3.1% by 21 days after the AAV-Cre injection (*p* = 0.027). We observed a 47.8 ± 4.79% increase in the protein levels of cleaved caspase 8 in the brains injected with AAV-Cre when compared to the AAV-GFP control, 21 days after injection (*p* = 0.04) and a striking 10-fold increase in the levels of cleaved caspase 3 (*p* = 0.014), 21 days after injection with AAV-Cre along with a significant increase in the expression of cleaved PARP 14 days after injection (*p* = 0.009). Thus, removal of KCC2 is also sufficient to induce apoptosis *in vivo.*

**Figure 6.**
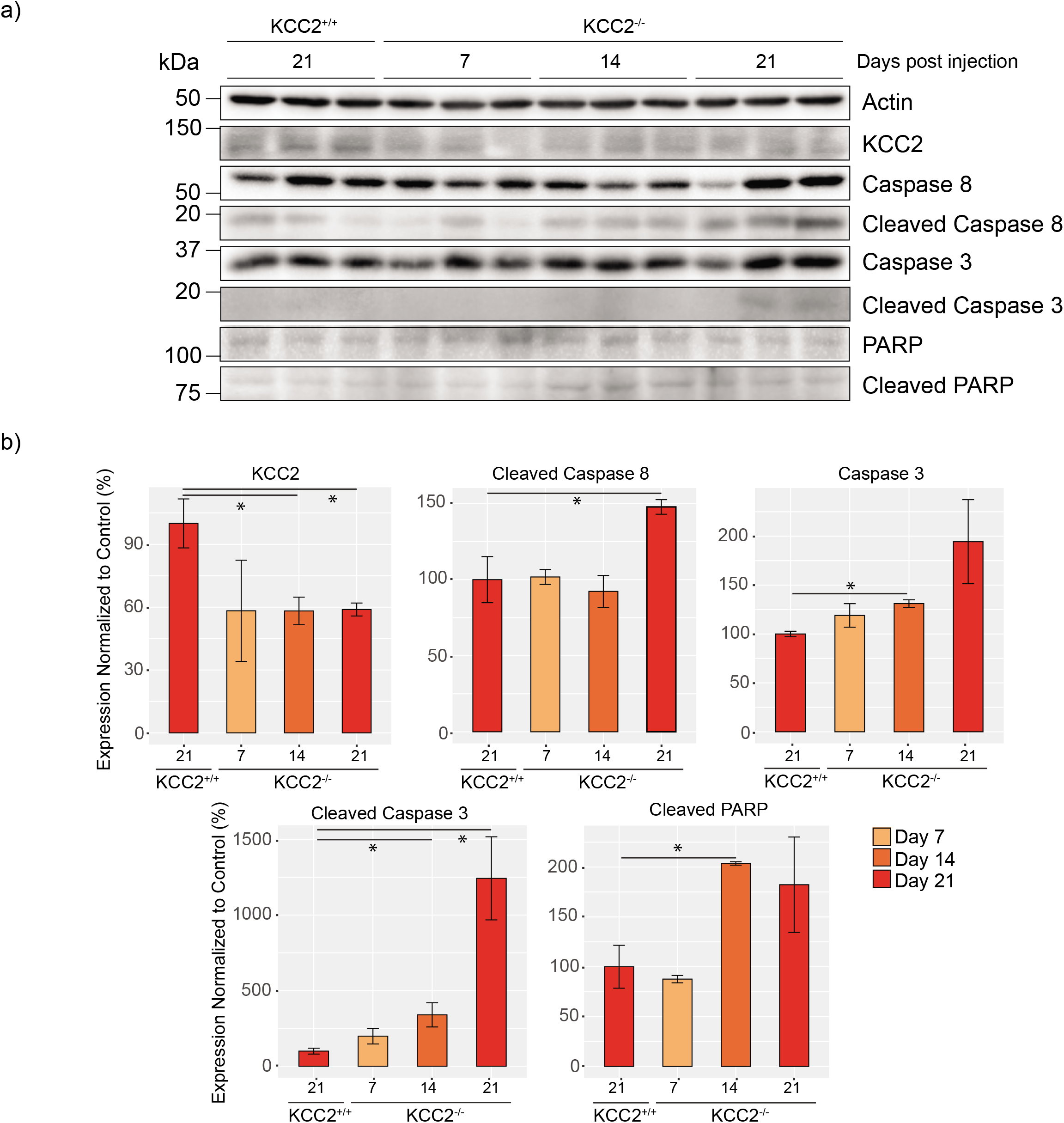
Loss of KCC2 *in vivo* increases the levels of proapoptotic markers. **A.** Immunoblots of hippocampal brain lysates harvested 7, 14 and 21 days after viral infection with AAV-Control or AAV-Cre probed for the total and cleaved levels of the extrinsic proapoptotic marker caspase 8, the intrinsic proapoptotic markers caspase 3 and caspase 7, the DNA damage marker PARP, KCC2, and β-actin. **B.** The bar chart shows the quantification of the levels of proapoptotic markers cleaved caspase 8, cleaved caspase 3 and cleaved PARP, with significantly changed expression marked (\**p* < 0.05).

## DISCUSSION

KCC2 provides the principle Cl^−^ extrusion mechanism for cells in the CNS. Here we used the previously described KCC2^fl^ mouse (16) combined with Cre-expressing AAVs to achieve highly efficient conditional removal of KCC2 in excitatory neurons (**Figure 1**). Using this strategy, we have begun to explore the role that KCC2 plays in determining neuronal morphology, neuronal survival, and the postnatal development of GABAergic inhibition in immature and mature neuronal networks. The placement of the *loxP* sites in the *SLC12A5* gene sequence of the KCC2^fl^ mouse, should result in a protein lacking the protein sequence flanked by Q911-Q1096. However, the data presented here compliments previous observations that truncation of the *SLC12A5* gene results in the ablation of KCC2 expression (15). This was demonstrated by immunoblotting with N- and C-terminal antibodies and suggests that either the truncated form of KCC2 protein is unstable and does not accumulate, or that the altered gene sequence results in premature transcriptional or translational termination. This is contrary to previous work (9, 10), and likely represents the differences associated with endogenous versus overexpression systems. We were not able to completely ablate KCC2 expression *in vitro* or *in vivo* under these experimental conditions. When KCC2 was removed from immature networks (DIV3 infected), that initially express low levels of KCC2, the residual KCC2 detected is likely expressed by interneurons (20). In mature networks however (DIV18 infected), where the KCC2 expression level is initially much higher, the residual KCC2 observed is likely to be a combination of interneuron expression and the result of slow protein turnover or local translation of slow turnover mRNAs. This is consistent with previous work estimating the half-life of KCC2 can be in the order of days (21).

Our approach aimed to differentiate between the removal of KCC2 in immature networks compared to mature networks, and study the resulting impact on neuronal morphology, activity and, viability *in vitro*. The methodology used in our *in vitro* experiments enables us to examine the effects of KCC2 removal with high temporal precision, that cannot be attained *in vivo,* as mice deficient in this transporter die shortly after birth (22). A recent study demonstrated that the genetic removal of KCC2 during neuronal migration induces the apoptosis of embryonic neocortical projection neurons (23). Yet, the functional consequences of KCC2 removal during postnatal neuronal development have not been investigated and are of great interest as a number of studies have suggested that KCC2 plays a transport independent role in determining spine morphology (9, 10). These studies used overexpression of a KCC2 construct devoid of its cytoplasmic tail (residues Q911-Q1096) or a DNA construct modified with a GFP reporter which may act to stabilize this truncated form of KCC2 (9, 10).

The ablation of KCC2 expression at DIV3 had no impact on the subsequent development of neurons (**Figure 2**). Both the complexity of proximal dendritic processes and spine formation were comparable to control cells. Likewise, ablating KCC2 expression in immature neurons did not induce cell death, as measured by cleaved caspases 3, 7, 8 or PARP, to detect apoptosis (**Figure 4**). However, hyperpolarizing GABA_A_R currents were compromised in neurons with reduced KCC2 expression (**Figure 3**) (15), despite relatively similar levels in inhibitory VGAT and gephyrin. Thus, the presence of KCC2 constitutes a prerequisite for GABAergic inhibition but may not contribute to neuronal morphological development or viability. Therefore, our results further highlight the critical role of KCC2 in the postnatal development of GABAergic inhibition, a process that has been proposed to be mediated in part by impermeant anions (24).

While the removal of KCC2 at DIV3 mimics the loss of KCC2 during development, the removal of KCC2 at DIV18, from mature networks which exhibit KCC2 activity-dependent hyperpolarizing GABA_A_R currents (1, 15), mimics the acute loss of KCC2 following ischemic injury or seizures (12, 25). Such acute neuronal injuries can cause gross neuronal damage, including alterations in spine morphology (26) and initiate the neuroinflammatory response, ultimately resulting in cell death (27). Following 6 days of viral exposure initiated at DIV18, we observed subtle changes in spine maturity consistent with previous observations (**Figure 2**) (9, 10). However, at this timepoint, we also detected multiple markers of apoptotic induction (**Figure 4**). Ablating KCC2 expression in mature cultures resulted in a significant increase in cleaved caspase 8 after 3 days of viral exposure. Caspase 8 is part of the extrinsic apoptotic pathway and is cleaved in response to extracellular signals. For instance, proteins such as tumor necrosis factor a (TNF-a) and tumor necrosis factor-related apoptosis-inducing ligand (TRAIL) bind to cognate death receptors, localized in the target cell plasma membrane (28). In the brain, these proteins are released in response to neuronal injury (29, 30). The cleavage of caspase 8 can result in both initiating the downstream caspase cascade leading to apoptosis, or to perpetuate the inflammatory response (31), and has been demonstrated in seizure-induced apoptotic cell death (32, 33). One of the immediate downstream consequences of caspase 8 cleavage is calpain activation, which in turn cleaves numerous protein targets. Sptan1 is a KCC2 associated protein (17) and is cleaved by calpains and caspases in response to neuronal injury (34) in a similar manner to KCC2 itself (21). After 6 days of viral exposure, loss of full length Sptan1 was observed, as well as cleaved downstream proteins; caspases 3, 7, and PARP. Cleavage of these proteins is indicative of the final stages of apoptotic cell death (35). These results were also mirrored in hippocampal tissue *in vivo*, and using the specific KCC2 inhibitor; 11K *in vitro* (36), which rapidly induced caspase 8, 3, and PARP cleavage within 30 minutes. Taken together, these results indicate that removal of KCC2 from mature neuronal networks causes changes in spine morphology, however this may at least be due to the induction of apoptosis via the extrinsic pathway. Data presented here suggests that changes in spine morphology, following KCC2 removal, are due to an apoptotic event and impact the notion that KCC2 has transporter independent functions.

Here, we have developed an elegant method of studying immature and mature neuronal networks in the presence or absence of KCC2. Our results demonstrate that reducing KCC2 expression levels in immature neurons does not impact gross neuronal morphology or viability but prevents the development of GABAergic inhibition. In contrast, mature neuronal networks are highly sensitive to KCC2 removal, where a reduction of approximately 50% is sufficient to induce apoptosis. Whether this is a direct consequence of the absence of KCC2 or due to subsequent excitotoxicity remains unanswered. However, this work advances our understanding of the pathological mechanisms that proceed reduced KCC2 expression and suggests that restoring KCC2 activity following events such as seizures or ischemia may prevent neuronal loss by apoptosis.

## EXPERIMENTAL PROCEEDURES

### Animal Care

Animal studies were performed according to protocols approved by the Institutional Animal Care and Use Committee of Tufts Medical Center (IACUC). Mice were kept on a 12 h light/dark cycle with *ad libitum* access to food and water.

### Antibodies

The primary antibodies used in this study were: KCC2 (mouse, Neuromab 75-013, WB 1:1000, ICC 1:500-1000), KCC2 (rabbit, LSBio LS-C135150, WB 1:1000), KCC2 (rabbit, Millipore 07-432, ICC 1:500-1000), GFP (chicken, Abcam 13970, ICC 1:1000), Caspase 3 (rabbit, CST #9662, WB 1:1000), Cleaved Caspase 3 (rabbit mAb, CST #9664, WB 1:1000), Caspase 7 (rabbit mAb, CST #12827, WB 1:1000), Cleaved Caspase 7 (rabbit mAb, CST #8438, WB 1: 1000), Caspase 8 (rabbit mAb, CST #4790, WB 1:1000), Cleaved Caspase 8 (rabbit mAb, CST #8592, WB 1:1000), Caspase 9 (mouse, CST #9508, WB 1:1000), Cleaved Caspase 9 (rabbit mAb, CST #52873, WB 1:1000), Caspase 12 (rabbit, CST #2202, WB 1:1000), PARP (rabbit, CST #9542, WB 1:1000), Cleaved PARP (rabbit mAb, CST #5625, WB 1:1000), Sptan1 (mouse, Abcam 11755, WB 1:1000) and β-actin (mouse, Sigma A1978, WB 1:5000). The secondary antibodies used were; goat anti-chicken Alexa Fluor 488 (Thermofisher, 1:1000), goat anti-rabbit Alexa Fluor 555 (Thermofisher, 1:1000), goat antirabbit Alexa Fluor 568 (Thermofisher, 1:1000), goat anti-rabbit Alexa Fluor 647 (Thermofisher, 1:1000), donkey anti-rabbit conjugated HRP (Jackson ImmunoResearch, 1:5000) and donkey anti-mouse conjugated HRP (Jackson ImmunoResearch, 1:5000).

### Electrophysiology

Electrophysiological experiments were carried out as previously described (15). Briefly, neurons were recorded at 33°C in saline containing 140 mM NaCl, 2.5 mM KCl, 2.5 mM MgCl2, 2.5 mM CaCl2, 10 mM HEPES, 11 mM glucose, pH 7.4 (NaOH). All solutions were applied through three-barrel microperfusion apparatus (700 μm, Warner Instruments, Hampden, CT), and conducted in the presence of tetrodotoxin (500 nM) and bumetanide (10 μM) to block voltage-dependent sodium channels and NKCC1 respectively. Cells infected with AAV9-CaMKII-eGFP (AAV-GFP), or AAV9-CaMKII-eGFP-Cre (AAV-Cre) (University of Pennsylvania Vector Core, Philadelphia, PA) were identified based on GFP expression using an epifluorescence microscope. Data were acquired using Clampex 10 software at an acquisition rate of 10 kHz with an Axopatch 2B amplifier and digitized with a Digidata 1440 (Molecular Devices, San Jose, CA). To detect endogenous Cl^−^ levels, cells were perforated with gramicidin (50 mg/mL) dissolved in an internal pipette containing 140 mM KCl, 10 mM HEPES, pH 7.4 (KOH) and a tip resistance of 3-4 MΩ. Following perforation to below a series resistance of 100 MΩ, E_GABA_ values were detected by application of muscimol (1 μM) during positive-going voltage ramps (20 mV/s).

### Generation and Genotyping of KCC2^fl^ Mice

KCC2^f1^ mice (*Slc12a5lox/lox*) were described previously (15, 16) and have been backcrossed on the C57BL/6 J background for at least 10 generations.

### Image Analysis

For immunoblots, individual band intensity was quantified using densitometry on Fiji, normalized to β-actin and further normalized to the corresponding control condition where appropriate. For immunocytochemistry (ICC), the KCC2 fluorescent intensity in 1024 x 1024 confocal images was quantified on Fiji and normalized to the control (37). Briefly, a mask was generated manually or using the thresholded GFP signal, that was then superimposed on the KCC2 fluorescent channel to measure the signal within the GFP positive region. The GFP signal was also used to reconstruct proximal dendrites manually, generate a .SWC file and perform the Sholl analysis using the Simple Neurite Tracer Plugin on Fiji (38).

### Immunoblotting

Immunoblotting was carried out as previously described (30). For cultured neurons, media was removed and spun down at 1000g for 5 minutes to harvest floating cells. Adherent cells were washed twice with phosphate buffered saline (PBS), combined with floating cells and lysed in RIPA buffer containing proteases and phosphatase inhibitors. For wild type cultured neurons treated with 11K, cells were washed in PBS and lysed in RIPA buffer containing proteases and phosphatase inhibitors. For tissue, AAV-GFP or AAV-Cre infected hippocampi were microdissected under a fluorescent microscope. The tissue was homogenized in RIPA buffer containing protease and phosphatase inhibitors. Proteins were quantified by Bradford assay. Samples were diluted to the same concentration in RIPA buffer and 2x sample buffer added. Samples were boiled for 5 mins at 95°C before being loaded onto polyacrylamide gels for sodium dodecyl sulphate poly acrylamide gel electrophoresis (SDS-PAGE). Proteins were then transferred onto nitrocellulose membranes, blocked in milk for 1 hour, and probed with primary antibodies overnight. The membranes were washed and probed with appropriate HRP-conjugated secondary antibodies and developed with enhanced chemiluminescent (ECL) substrate. Where possible all replicates were run on the same gel.

### Immunocytochemistry

Immunocytochemistry (ICC) was carried out as previously described (17). Briefly, primary cultured hippocampal/cortical neurons were washed with PBS, fixed in 4% paraformaldehyde (PFA, Electron Microscopy Services, Hatfield, PA) in PBS, and permeabilized in block solution consisting of 1X PBS with 0.1% Triton X-100, 3% (w/v) Bovine Serum Albumin (BSA), 10% normal goat serum and 0.2 M glycine. Primary and secondary antibodies (as described above) were prepared at a dilution of 1:1000 in block solution and incubated with the cells for 1 h at room temperature in the dark. The cells were washed in 1X PBS and mounted on glass slides using ProLong Gold.

### Primary Neuronal Culture

Primary cortical/hippocampal neurons were prepared and cultured as previously described (39). Briefly, P0 KCC2^fl^ mice were anesthetized on ice and the brains removed. The brains were dissected in Hank’s buffered salt solution (HBSS, Invitrogen) with 10 mM HEPES. The cortices and hippocampi were trypsinized and triturated to dissociate the neurons. Cells were counted using a hemocytometer and plated on poly-L-lysine-coated coverslips (for ICC and electrophysiology) or in 35 mm dishes (for immunoblot) at a density of 1 × 10^5^ or 4 × 10^5^ cells respectively. At days *in vitro* (DIV) 3 or 18, cells were exposed to AAV-GFP or AAV-Cre as described above at a concentration of 1 × 10^6^ genome copies (GC)/mL. For biochemical analysis and ICC, cells infected at DIV3 were harvested at DIV7, 14 and 21 and cells infected at DIV18 were harvested at DIV18, DIV21 and DIV24. At DIV18-21, primary neurons were treated with either DMSO or 11K (1 μM) for 10, 30 and 60 minutes before harvested for biochemical analysis.

### Statistics

Fluorescent imaging and perforated patch-clamp data were subjected to the D’Agostino-Pearson omnibus normality test and the F-test to compare variances on Prism. To assess statistical significance, the unpaired two-sample T-test was performed, and the Welch’s correction was applied when necessary. The cumulative distributions generated from Sholl analysis were subjected to the Kolmogorov-Smirnov (KS) test. Significance of *p* < 0.05 is represented as *, *p* < 0.01 is represented as **, *p* < 0.001 is represented as *** and *p* < 0.0001 is represented as ****.

### Stereotaxic Injection

Age-matched male WT and KCC2^fl^ animals were bilaterally injected with adeno-associated virus (AAV) expressing Cre recombinase. Prior to surgery, animals were anaesthetized by isoflurane inhalation and administered buprenorphine HCl (0.1mg/kg) subcutaneously as analgesic. The deeply anaesthetized animals were immobilized and stabilized in a Kopf stereotaxic apparatus. Holes were drilled on the skull surface and Neurosyringe (Hamilton, Reno, NV) was used to deliver 1 μL of AAV-GFP or AAV-Cre (1 x 10^13^ GC/mL) into each of the CA1 areas with the following coordinates relative to Bregma: AP: −2.0, ML: +/-1.8, DV: −2.0. AAV was injected at a rate of 100nL/min, under the control of a microsyringe pump controller (World Precision Instrument Micro4). After the injection, the incision site was sutured, and animals were housed individually and postoperatively monitored.

### Tissue Processing

Animals were anaesthetized with isoflurane. Once the animals reached deep anesthesia state, they were decapitated, the brains were extracted and separated into two hemispheres. One hemisphere was processed for western blot analysis (See Immunoblotting) and the other was immersed in 4% PFA at 4°C overnight. The PFA-fixed brain hemispheres were kept in 30% sucrose solution at 4°C for cryoprotection for 3 days and embedded and frozen in Tissue-Tek® OCT-compound (VWR). 20 μm coronal sections were prepared with a cryostat (Leica HM525, Leica Microsystems, Wetzlar, Germany) and were stored at −20°C in cryoprotectant solution.

## Acknowledgements

S.J.M. is supported by National Institutes of Health (NIH)–National Institute of Neurological Disorders and Stroke Grants NS051195, NS056359, NS081735, R21NS080064 and NS087662; NIH–National Institute of Mental Health Grant MH097446. GK and JN are fellows of and funded by the AstraZeneca Postdoctoral Program.

## Author contributions

G.K performed experiments, analyzed data and wrote the paper. S.J.M and J.L.S conceptualized the project, designed experiments and wrote the paper. J.N, R.A.C, J.H, C.C, Q.R and J.L.S performed experiments. G.K, J.T.K, N.J.B, S.J.M and J.L.S edited the paper.

## Abbreviations

AAV: Adeno-associated virus
DIV: days *in vitro*
GABA_A_R: γ-aminobutyric acid type A receptors
GFP: green fluorescent protein
KCC2: K^+^/Cl^−^ co-transporter 2
KO: knock-out
PARP: poly (ADP-ribose) polymerase (PARP)
TLE: temporal lobe epilepsy

